# LOXL2 Mediates Airway Smooth Muscle Cell Matrix Stiffness and Drives Asthmatic Airway Remodelling

**DOI:** 10.1101/2020.11.16.384792

**Authors:** Jopeth Ramis, Robert Middlewick, Francesco Pappalardo, Jennifer T. Cairns, Iain D. Stewart, Alison E. John, Shams-un Nisa Naveed, Ramaswamy Krishnan, Suzanne Miller, Dominick E. Shaw, Christopher E. Brightling, Lee Buttery, Felicity Rose, Gisli Jenkins, Simon R. Johnson, Amanda L Tatler

## Abstract

Airway smooth muscle cells (ASM) are fundamental to asthma pathogenesis, influencing bronchoconstriction, airway hyper-responsiveness, and airway remodelling. Extracellular matrix (ECM) can influence tissue remodelling pathways, however, to date no study has investigated the effect of ASM ECM stiffness and crosslinking on the development of asthmatic airway remodelling. We hypothesised that TGFβ activation by ASM is influenced by ECM in asthma and sought to investigate the mechanisms involved.

This study combines in vitro and in vivo approaches: human ASM cells were used in vitro to investigate basal TGFβ activation and expression of ECM crosslinking enzymes. Human bronchial biopsies from asthmatic and non-asthmatic donors were used to confirm LOXL2 expression ASM. A chronic ovalbumin model of asthma was used to study the effect of LOXL2 inhibition on airway remodelling.

We found that ASM cells from asthmatics activated more TGFβ basally than non-asthmatic controls and that diseased cell-derived ECM influences levels of TGFβ activated. Our data demonstrate that the ECM crosslinking enzyme LOXL2 is increased in asthmatic ASM cells and in bronchial biopsies. Crucially, we show that LOXL2 inhibition reduces ECM stiffness and TGFβ activation in vitro, and can reduce subepithelial collagen deposition and ASM thickness, two features of airway remodelling, in an ovalbumin mouse model of asthma.

These data are the first to highlight a role for LOXL2 in the development of asthmatic airway remodelling and suggest that LOXL2 inhibition warrants further investigation as a potential therapy to reduce remodelling of the airways in severe asthma.

## Introduction

Airway remodelling is a common feature in asthma and can include epithelial shedding, increased airway smooth muscle (ASM) mass and subepithelial fibrosis. The extracellular matrix (ECM) is emerging as a key modulator of inflammatory and remodelling events [1–3]. Proteins present within asthmatic ECM can influence the secretory profile of ASM cells and promote their proliferation [4, 5], and influence airway remodelling, hyper-responsiveness and inflammation [6]. The relative stiffness of ECM can also influence cell behaviour [7, 8]. In the lung increased ECM stiffness initiates epithelial-mesenchymal transition and causes fibroblasts to synthesise collagen [9, 10]. Furthermore, increased matrix stiffness can increase ASM cell proliferation and contractile force [11, 12]. Together these studies suggest that increased ECM stiffness may contribute to the development of airway remodelling.

ECM stiffness is influenced by crosslinking, the formation of covalent bonds between ECM proteins, which is catalysed by ECM crosslinking enzymes. Lysyl oxidase like-2 (LOXL2) belongs to the lysyl oxidase (LOX) family of enzymes and is responsible for crosslinking and stabilising collagen fibres. Increased LOXL2 expression is evident in various fibrotic diseases [13–17], and LOXL2 inhibition reduces fibrosis in multiple animal models of disease [17–20]. To date, there have been no studies investigating a role for LOXL2 in asthma pathogenesis or airway remodelling.

This study investigated a role for ECM alterations and LOXL2 in the development of asthmatic airway remodelling. We found that activation of TGFβ by ASM cells is influenced by ECM and that ASM cells isolated from asthmatic donors exhibit enhanced basal TGFβ activation and deposit a stiffer ECM. We show that LOXL2 expression is increased in asthma and that LOXL2 inhibition reduces ASM cell ECM stiffness and TGFβ activation in vitro and reduces airway remodelling in an in vivo asthma model.

## Materials and Methods

### Cell culture

The source of human ASM cells is detailed in supplemental methods. ASM cells were cultured as previously described [21] and growth arrested in serum-free DMEM for 24 hours prior to experiments. A minimum of 3 donor cell lines were used in each experiment.

### Culture of ASM Cells on Methacrylated Gelatin (GelMa) Hydrogels

A hydrogel disc was produced from a precursor gelatin methacrylol (GelMA) solution containing 0.5% Irgacure 2959 by crosslinking under UV light (365nm). Solutions of 5, 10 and 15% wt/vol GelMA were dissolved to obtain hydrogels that were 1x, 6x, and 12x that of the physiological stiffness of tracheal smooth muscle reported previously [22]. Hydrogel relative stiffness (E) was determined by texture analyser (Supplemental Figure S1). ASM cells were seeded at 2,000 cells/cm^2^ and cultured for 6 days.

### Phosphorylated Smad2 (PSmad2) Immunofluorescence

Cells were fixed (4% paraformaldehyde), permeabilised (0.3% Triton-X100) then stained with PSmad2 primary antibody (1:100) and an Alexa Fluor™-488 antibody (1:3000). Nuclei were stained with DAPI (4’,6-Diamidino-2-Phenylindole, Dihydrochloride). Image analysis was performed using a Zeiss LSM 710 confocal microscope and Cell Profiler™[23].

### Cyclical Stretching

ASM cells were seeded fully confluent on collagen I coated BioFlex^®^ plates and 15% stretch (0.3Hz) was applied using the Flexcell^®^ FX-6000T Tension system. Additional plates were incubated alongside the Flexcell^®^ device to serve as unstretched controls.

### Western Blotting

PSmad2 western blotting was performed as previously described [24]. To measure LOXL2 and LOXL3 expression cell lysates were electrophoresed, transferred on to PVDF membrane then probed using antibodies against LOXL2 (1:1000), LOXL3 (1:2000) and GAPDH (1:3000). Bands were visualised using Clarity ECL and densitometry was performed using Image J.

### Transformed Mink Lung Epithelial Cell (TMLC) Assay

TMLC coculture assay was performed as previously described [24] using ASM cells seeded in to 96 well plates at 4.025 × 10^4^ cells/ well.

### PSmad2 ELISA

PSmad2 in cell lysates was measured using an ELISA kit according to the manufacturer’s instructions. Due to the absence of a standard for generation of a standard curve data are presented as raw optical density (OD) values.

### Collagen Gel Contraction Assay

Contractility was assessed using a cell contraction assay according to the manufacturer’s instructions. Briefly, ASM cells were seeded at 0.5×10^6^ cells/well of 24 well plates and the gel diameter was quantified after 24 hours using a Nikon SMZ1500 stereomicroscope and Image J. Positive and negative controls were performed using methacholine and 2,3-butanedione monoxime, respectively.

### Traction Force Microscopy (TFM)

ASM cells were seeded at 2×10^4^ cells/well onto 3kPa polydimethylsiloxane (PDMS) NuSil gel-8100 coated 96-well plates with embedded fluorescent microspheres, [25]. Fluorescent bead displacement was used to calculate traction forces using the method of Fourier Transform traction cytometry [26]. Traction force data was obtained from 8-24 separate wells per cell line. The root mean square (RMS) value for traction in Pascals (Pa) was reported.

### ECM Cross-over Experiments

ASM cells were seeded at 4.025×10^5^ cells/ml and cultured for three days then digested in 0.016M NH4OH for 30 minutes. Non-asthmatic cells were seeded on to either their own ECM or on to ECM deposited by asthmatic ASM cells at 4.025 × 10^5^ cells/ml, and vice versa for asthmatic ASM cells. End-point analysis was performed 24 hours later.

### Quantitative RT-PCR (qRT-PCR)

Gene expression changes were assessed using qRT-PCR as previously described [27]. Primer sequences are shown in Supplemental Table 2.

### Atomic Force Microscopy (AFM)

The MFP-3D Standalone Atomic Force Microscope was used to obtain Force-displacement (F-D) curves of ECM for Young’s modulus (E) calculation. Instrument sensitivity was calibrated with the unloading F-D curve slope on glass, whilst the instrument thermal fluctuations were used to extract the effective spring constant of the tip, inherent to its resonant frequency. For each measurement of the sample, at least 100 F-D curves were recorded.

### Bronchial Biopsy Collection

Bronchial biopsies obtained from asthmatic and healthy control donors (n=6/group) were used to determine LOXL2 expression within ASM bundles. All tissue was collected at the Nottingham Biomedical Research Centre, University of Nottingham, under informed, written consent with ethical approval (12/EM/0199). More details provided in supplemental methods.

### Immunohistochemistry

Parallel 5μm sections of lung tissue were boiled in citrate buffer then incubated with either a LOXL2 (1:1000) or αSMA (1:1000) antibody overnight followed by a goat anti-rabbit antibody (1:200). The sections were incubated with 3,3’-diaminobenzidine and counterstained with Mayer’s haematoxylin. ImageJ was used to overlay positive LOXL2 staining on αSMA regions to quantify percentage reactivity.

### LOXL2 Inhibition in an In Vivo Ovalbumin Model

Studies were approved by the University of Nottingham Animal and Welfare Ethical Review Board and performed under Home Office license authority within the Animal (Scientific Procedures) Act 1986. Following sensitisation by intraperitoneal injection of ovalbumin/ALUM, animals were randomised to one of four treatment groups (PBS + vehicle, PBS + PAT1251, OVA + vehicle, OVA + PAT1251). Full details of the model provided in supplemental methods.

### Histology

Haematoxylin and eosin and Masson’s trichrome staining of 5μm thick lung sections was performed as previously described [28, 29]. Staining was visualised using a Nikon Eclipse 90i microscope and NIS Elements AR3.2 software (Nikon).

### Immunostaining

Murine lung tissue was subjected to αSMA immunofluorescence and quantification of small airway (radius less than 100μm) staining was performed as previously described [21].

### Statistical Analysis

All data are reported as a median of n observations and non-parametric tests were used in all cases. All statistical tests were discussed with a statistician (I.D.S). Further details are included in the supplemental methods. Details of the statistical test used for each figure is included in the figure legend.

## Results

### Substrate Stiffness and Cyclical Stretch Activates TGFβ in Human ASM Cells

We have previously demonstrated that contractile agonists increase integrin-mediated TGFβ activation in human ASM cells [21]. Since both substrate stiffness and cyclical stretch can influence integrin-mediated TGFβ activation in epithelial cells [30, 31] we investigated whether these processes affect TGFβ activation in ASM cells. Human ASM cells were cultured on GelMa hydrogels of increasing stiffness for seven days. The stiffness of the hydrogels was determined using texture analyser (Supplemental Figure S1). Active TGFβ signalling (nuclear PSmad2) was present when human ASM cells were cultured on a substrate of equivalent stiffness to the physiological stiffness of tracheal smooth muscle (1x) [22] and levels increased with increased substrate stiffness (Figure 1A and 1B). Additionally, cyclical mechanical stretch caused increased PSmad2 levels after 4 hours (Figure 1C and 1D). Together, these data show that basal TGFβ activation by ASM cells can be influenced by both substrate stiffness and cyclical stretching.

**Figure 1.**
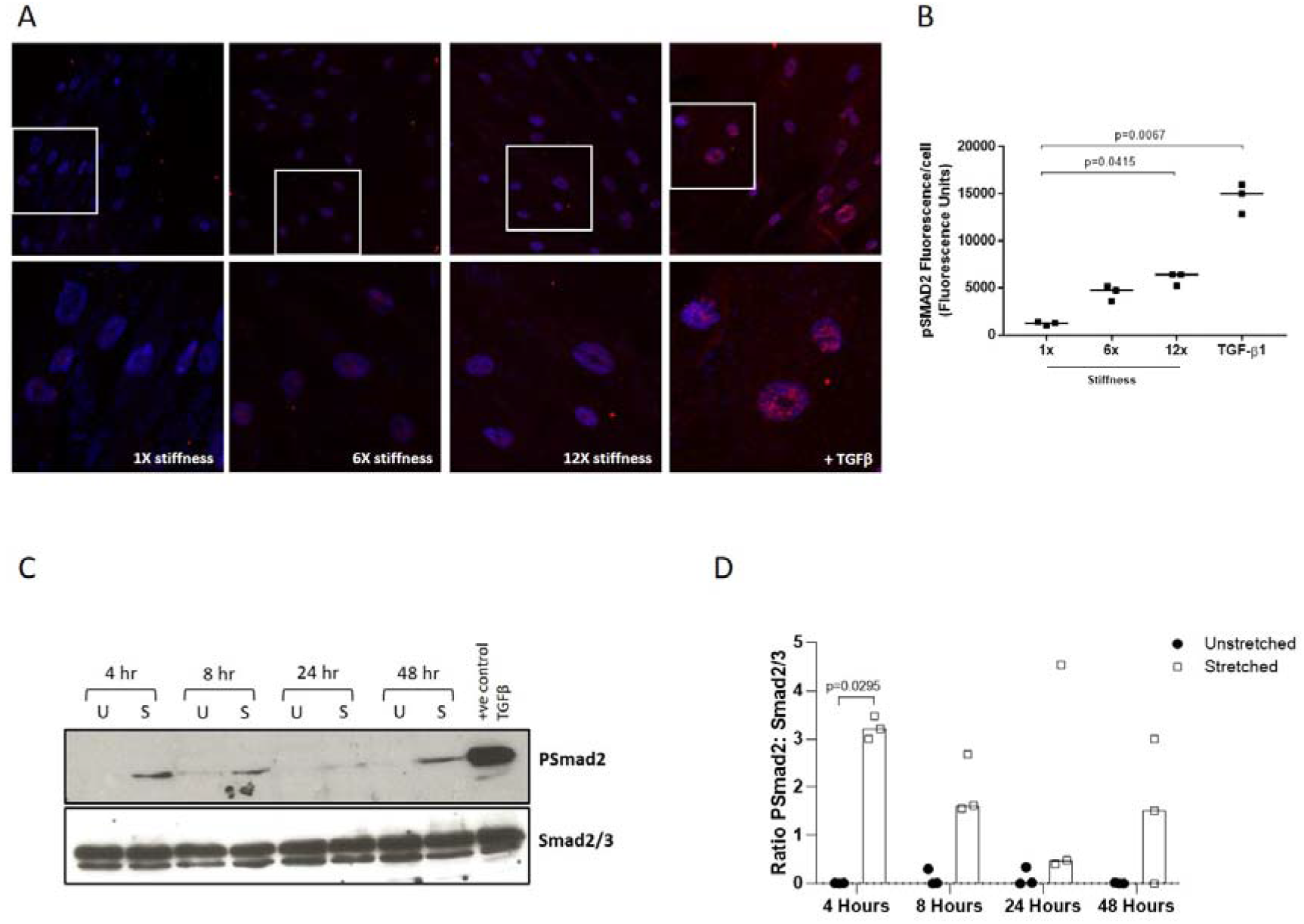
A Representative immunofluorescence images of non-asthmatic ASM cells cultured on GelMa substrates of increasing stiffness (1x, 6x and 12x that of physiological stiffness [22]). TGFβ (5ng/ml) was used as a positive control. The cells were stained for PSmad2 (Alexa Fluor™ 488) and DAPI and imaged using confocal microscopy at 20x magnification. B Nuclear PSmad2 immunofluorescence was quantified in non-asthmatic ASM cells cultured on GelMa substrates of increasing stiffness (1x, 6x and 12x), or on glass coverslips and treated with TGFβ (5ng/ml) as a positive control. Data is presented as median fluorescence units from three individual donor cell lines. The number of cells analysed was 106-305/donor/condition and the experiment was independently repeatedly twice. Statistical analysis was performed using Kruskall Wallis non-parametric test with Dunn’s multiple comparison test. C Non-asthmatic ASM cells were subjected to 15% stretch at 0.3Hz (S) or left unstretched (U) for 4, 8, 24 and 48 hours. PSmad2 and total Smad2/3 were measured by western blotting. Figure shown is representative of n=3 donor cells. D Densitometrical analysis of the western blots outlined in Figure 1C was performed using Image J and the data are shown here as median ratio of PSmad2: Smad2/3. Statistical analysis was performed using Kruskall Wallis non-parametric test with Dunn’s multiple comparison test

### Asthmatic Airway Smooth Muscle Cells Activate Increased Levels of TGFβ

We next sought to investigate whether basal TGFβ activation by ASM cells is aberrant in asthma. Basal TGFβ activation was increased in ASM cells isolated from asthmatic donors compared with non-asthmatic cells as measured by a TMLC co-culture assay (Figure 2A) and by PSmad2 ELISA (Figure 2B). As we, and others, have shown a role for cell contractility in regulating TGFβ activation, we measured cell contractility. Contractility of asthmatic ASM cells was increased compared with non-asthmatic controls when measured using a collagen gel contraction assay (Figure 2C) and the degree of contractility correlated with the amount of TGFβ activated by the cells (Figure 2D). However, cell contraction was similar between asthmatic and non-asthmatic ASM cells when measured by traction force microscopy (Supplemental Figure S2A).

**Figure 2.**
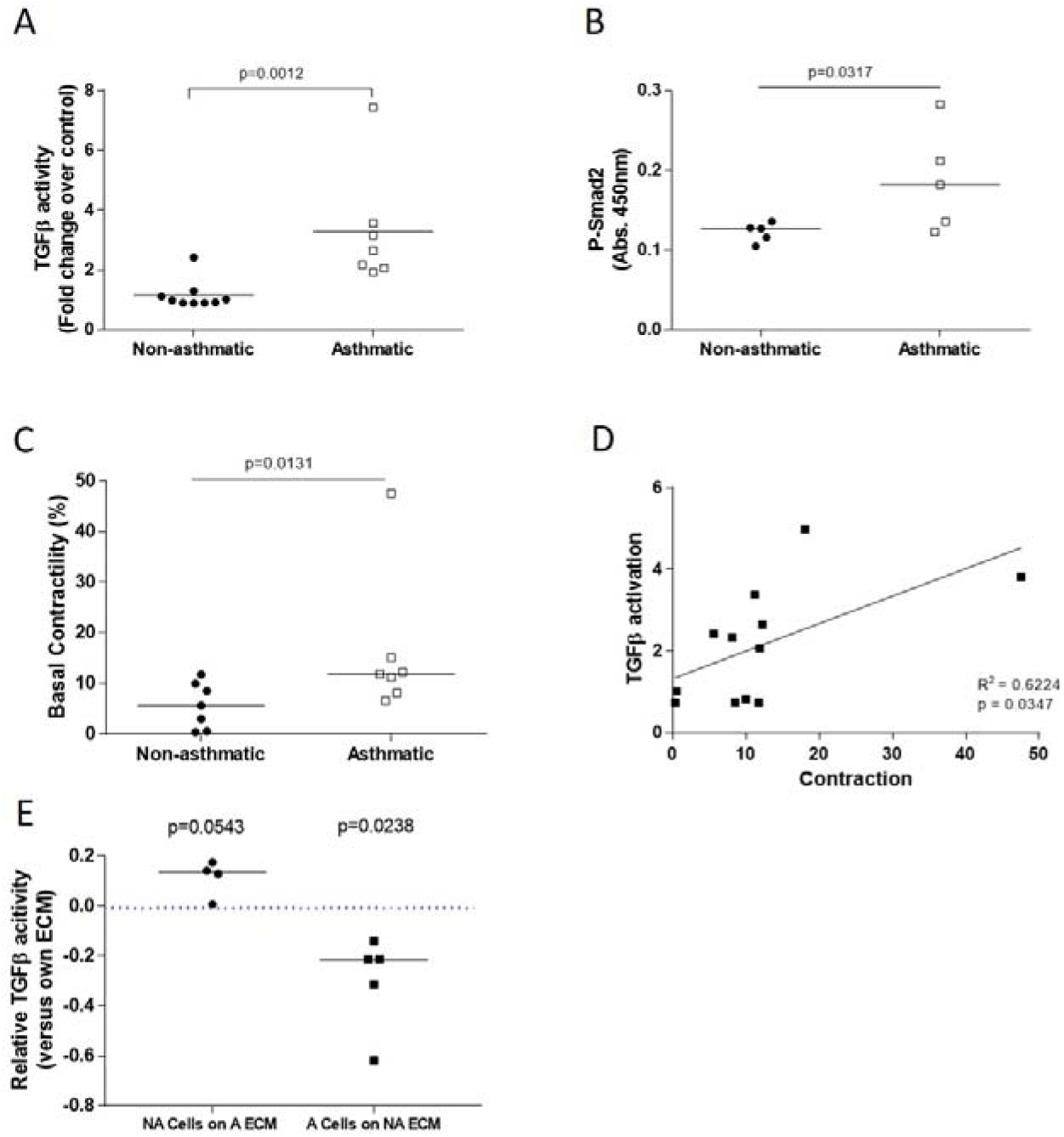
A Basal TGFβ activity in non-asthmatic and asthmatic ASM cells was measured by TMLC reporter cell assay. Data is expressed as median fold change in TGFβ activity in pg/ml per 10^4^ cells versus the mean data for non-asthmatic cells. Statistical analysis was performed using non-parametric Mann Whitney test. B Basal TGFβ activation in non-asthmatic and asthmatic ASM cells was measured using a PSmad2 ELISA. Data is expressed as median optical density (450nm). Statistical analysis was performed using non-parametric Mann Whitney test. C Basal cell contractility in non-asthmatic and asthmatic ASM cells was measured using a collagen gel contraction assay. Data is expressed as median percentage contraction of the collagen gel. Statistical analysis was performed using non-parametric Mann Whitney test. D Basal TGFβ activity (Figure 2A) was correlated with basal cell contractility using a Spearman correlation and the R^2^ value calculated (Figure 2C). E ECM crossover experiments were performed where non-athmatic ASM cells were cultures on asthmatic ECM and vice versa prior to determination of TGFb activity by TMLC assay. Data for each individual donor cell line is expressed relative to TGFβ activity levels when cultured on the cell’s own ECM. The blue dotted line denotes no change in TGFβ activity when cultured on different ECM. Both asthmatic and non-asthmatic data is presented on a single graph to illustrate the direction of change in TGFβ activity when cultured on the opposing ECM. Data is expressed as median relative TGFβ activity. Statistical analysis was performed using a one-sample T Test.

To further explore whether differences in cytoskeleton-mediated TGFβ activation may explain the observed phenotype of enhanced TGFβ activation in asthmatic ASM cells we applied cyclical mechanical stretch. Stretching caused increased PSmad2 in both asthmatic and non-asthmatic ASM cells (Supplemental Figure S2B and C). We found no significant difference in PSmad2 levels in stretched asthmatic ASMs compared with non-asthmatic ASMs (Supplemental Figure S2B and C). We concluded that while contraction and cytoskeletal alterations affect basal TGFβ activation it was unlikely to be solely responsible for driving enhanced TGFβ activation in asthmatic ASM cells.

We hypothesised that ECM alterations could be an additional factor driving increased TGFβ activation in asthmatic ASM cells. We performed ECM crossover experiments using decellularised ECM preparations. Culturing non-asthmatic ASM cells on asthmatic cell-derived ECM increased TGFβ activation compared with culturing on their own endogenous ECM (Figure 2E). Similarly, culturing asthmatic ASM cells on non-asthmatic ECM led to a significant decrease in TGFβ activation (Figure 2E).

### The ECM Crosslinking Enzyme LOXL2 Is Increased in Asthma and Regulates ECM Stiffness

To investigate a mechanism driving alterations in asthmatic ECM we assessed expression of ECM crosslinking enzymes. LOXL2 and LOXL3 mRNA were significantly increased in asthmatic ASM cells compared with controls (Figure 3A and B). There was also a trend towards increased TGM2 and LOX expression (Supplemental Figure S3A and B). We sought to confirm increased LOXL2 and LOXL3 at the protein level (Figure 3C). LOXL2 protein expression was increased in asthmatic ASM cells compared with non-asthmatic cells (Figures 3C and D). Despite increased levels of LOXL3 mRNA (Figure 3B) expression of LOXL3 protein was reduced in asthmatic ASM cells (Figure 3C and Supplemental Figure S3C).

**Figure 3.**
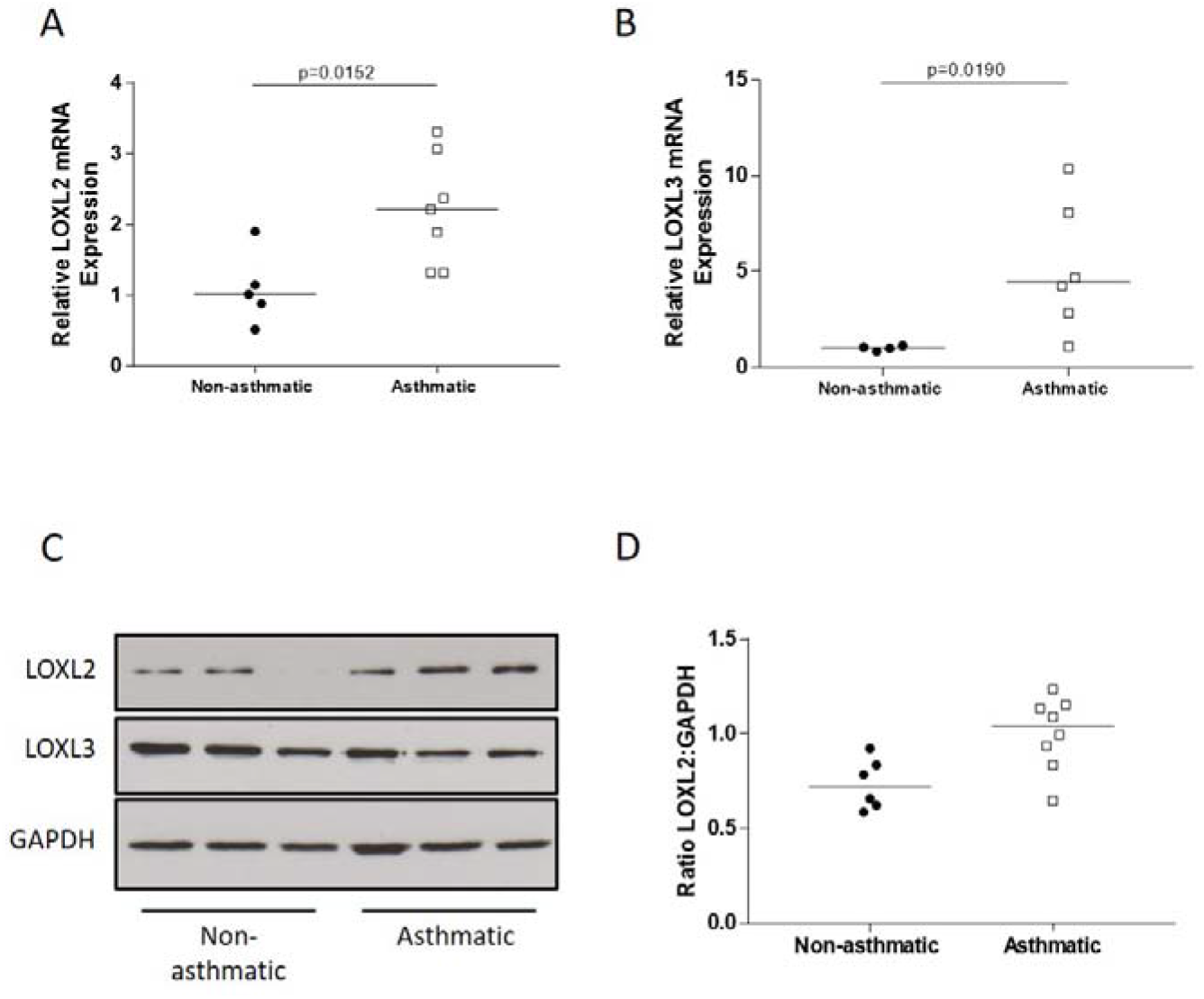
A Relative *LOXL2* mRNA in non-asthmatic and asthmatic ASM cells was determined by qRT-PCR. Data is expressed as median fold change in *LOXL2* mRNA versus the mean data from the non-asthmatic group. Statistical analysis performed by non-parametric Mann Whitney test. B Relative *LOXL3* mRNA in non-asthmatic and asthmatic ASM cells was determined by qRT-PCR. Data is expressed as median fold change in *LOXL3* mRNA versus the mean data from the non-asthmatic group. Statistical analysis performed by non-parametric Mann Whitney test. C Representative western blots for LOXL2, LOXL3 and GAPDH in non-asthmatic (n=3) and asthmatic (n=3) ASM cells. D Densitometrical analysis of western blots for LOXL2 in non-asthmatic and asthmatic ASM cells. Data is expressed as median ratio LOXL2 to GAPDH from two separate western blots (total n=6 non-asthmatic and n=8 asthmatic donor cell lines). Statistical analysis was performed by non-parametric Mann Whitney test.

To further investigate whether LOXL2 expression is increased in asthma we stained parallel sections of airway biopsies from non-asthmatic and asthmatic donors for both α-SMA and LOXL2 (Figure 4A). We quantified the percentage of smooth muscle (α-SMA positive) within the biopsy that was positive for LOXL2 (Figure 4B). Our data demonstrated variability in LOXL2 positivity within smooth muscle bundles of asthmatic biopsies, including high levels that were not observed in non-asthmatic controls (Figure 4B).

**Figure 4.**
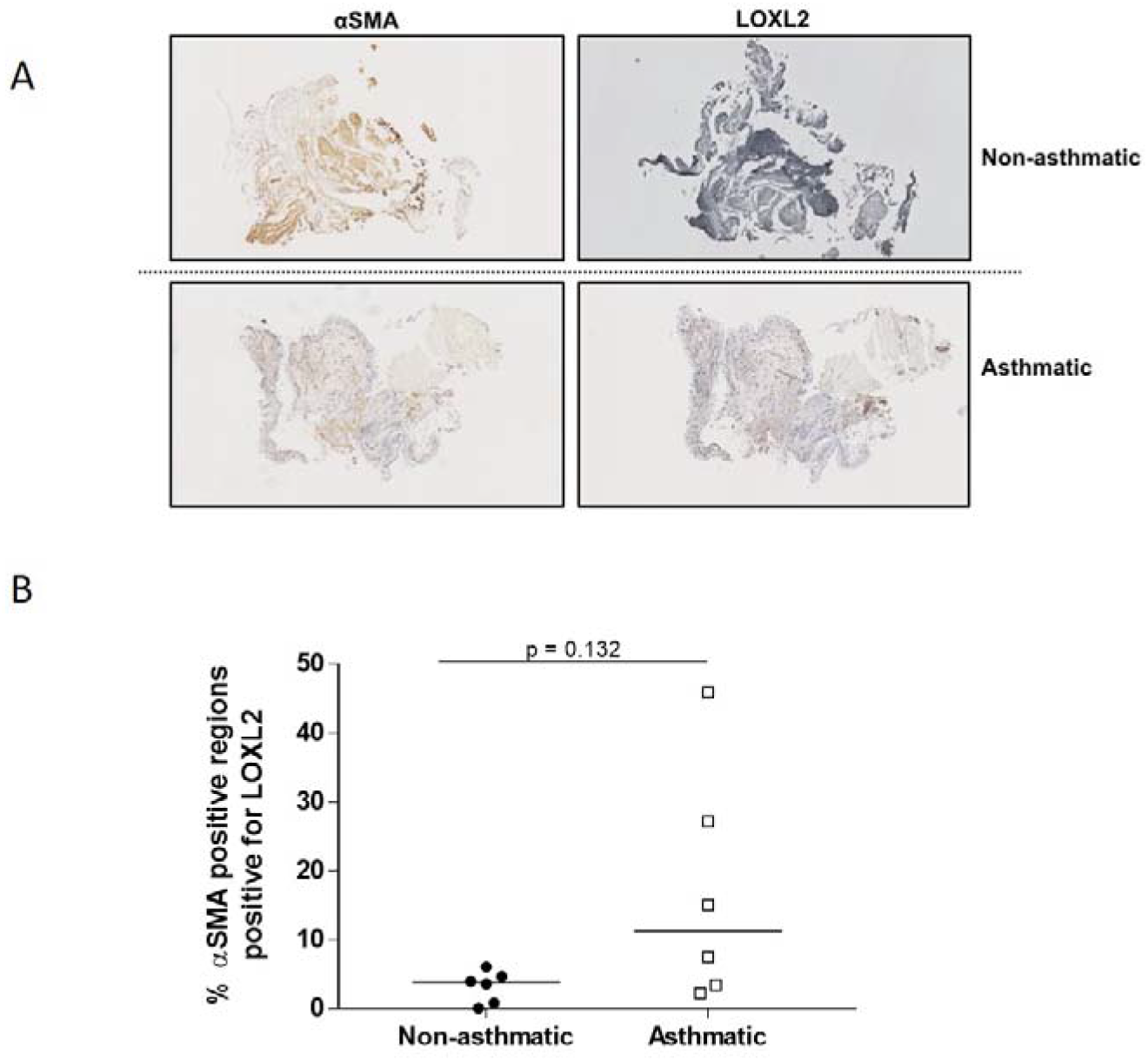
A Representative images demonstrating αSMA and LOXL2 immunostaining in parallel sections from biopsies from a non-asthmatic and an asthmatic donor. Images of immunostained biopsies were imaged using NanoZoom^®^. Figures show the resulting images at 5x magnification. B Quantification of LOXL2 staining in asthmatic and non-asthmatic human bronchial biopsies described above (Figure 4C). Image J was used to identify the αSMA positive regions within the biopsy then overlay the LOXL2 image to allow quantification of the percentage of αSMA positive regions that were positive for LOXL2. Data is expressed as median percentage of αSMA positive regions that were positive for LOXL2 staining. Statistical analysis was performed using non-parametric Mann Whitney test.

### LOXL2 Inhibition Reduces ECM Stiffness and Allergen-Induced Airway Remodelling

To investigate the effect of LOXL2 on the stiffness of ECM deposited by ASM cells we treated ASM cells with a LOXL2 inhibitor (PAT1251; IC50 0.71μM). 1μM PAT1251 reduced the stiffness of ECM deposited by asthmatic ASM cells to an equivalent stiffness as that deposited by non-asthmatic controls (Figure 5A). Furthermore, 1μM PAT1251 led to a small but significant reduction in basal TGFβ activation by asthmatic cells (Figure 5B).

**Figure 5.**
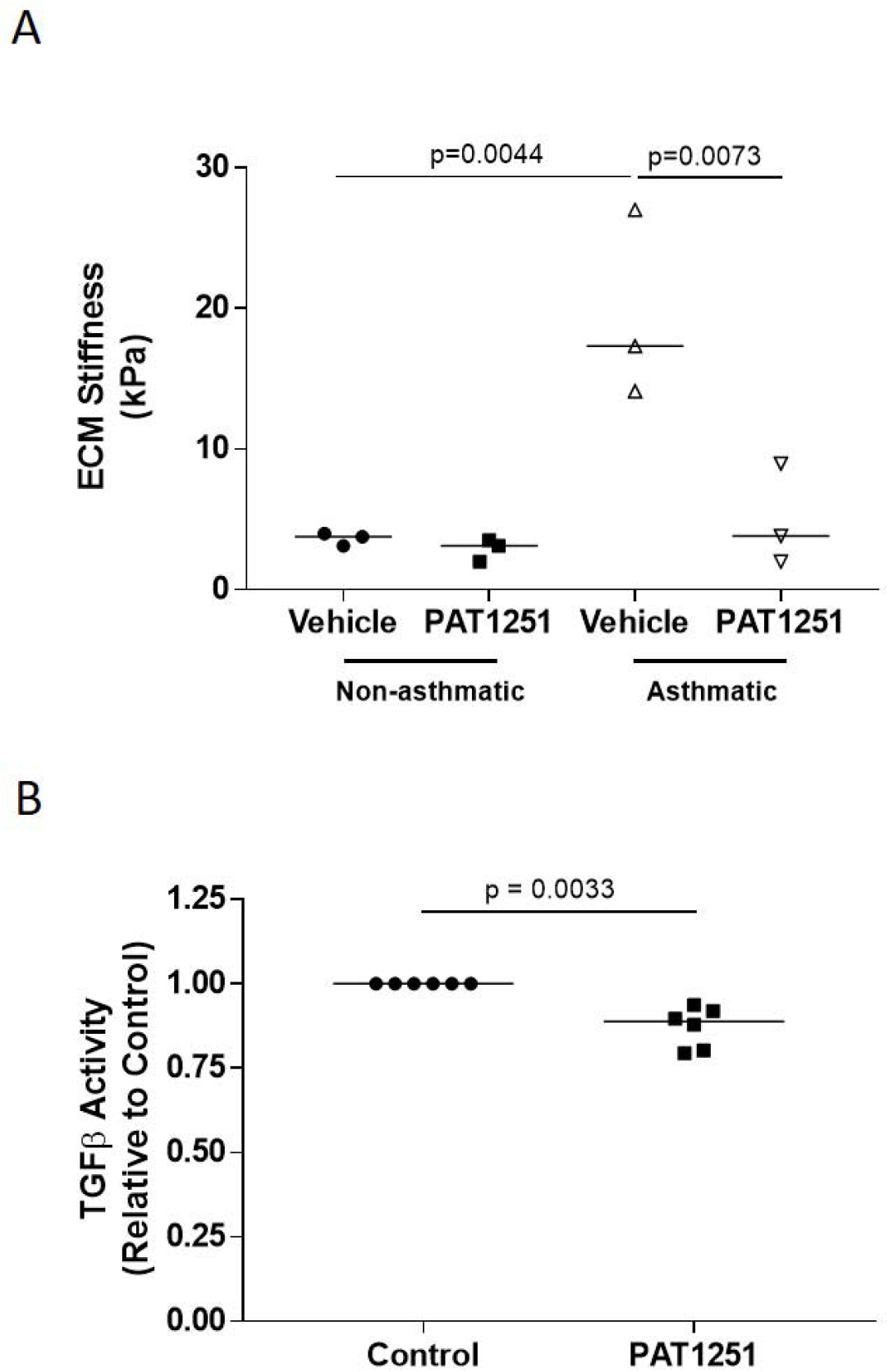
A ECM deposited by non-asthmatic and asthmatic ASM cells cultured in the presence of either DMSO vehicle control or LOXL2 inhibitor (PAT1251; 1μM) was subjected to AFM to measure the ECM stiffness. Data is expressed as median ECM stiffness (kPa). Statistical analysis was performed using Kruskall Wallis non-parametric test with Dunn’s multiple comparison test. B Confluent asthmatic ASM cells were cultured for three days in either DMSO vehicle control or LOXL2 inhibitor (PAT1251; 1μM) then TGFβ activity assessed by TMLC assay. Data are presented median TGFβ activity relative to vehicle control. Statistical analysis was performed using a one sample t-test.

We next investigated a role for LOXL2 in ovalbumin-induced airway remodelling in vivo. Daily oral dosing with 30mg/kg PAT1251 [18] from the start of ovalbumin challenges protected mice from ovalbumin challenge related decreases in body mass (Figure 6A). There was no significant difference in starting body mass between treatment groups (Supplemental Figure S4A). Ovalbumin challenge led to increased BAL inflammation (Figure 6B and C), including increased percentages of eosinophils, lymphocytes and neutrophils and a concomitant decrease in macrophages (Supplemental Figures S4B-E). PAT1251 had no significant effect on ovalbumin challenge-induced airway inflammation (Figure 6B and 6C), nor did it affect the differential cell counts of eosinophils, lymphocytes, neutrophils or macrophages (Supplemental Figures S4B-E).

**Figure 6.**
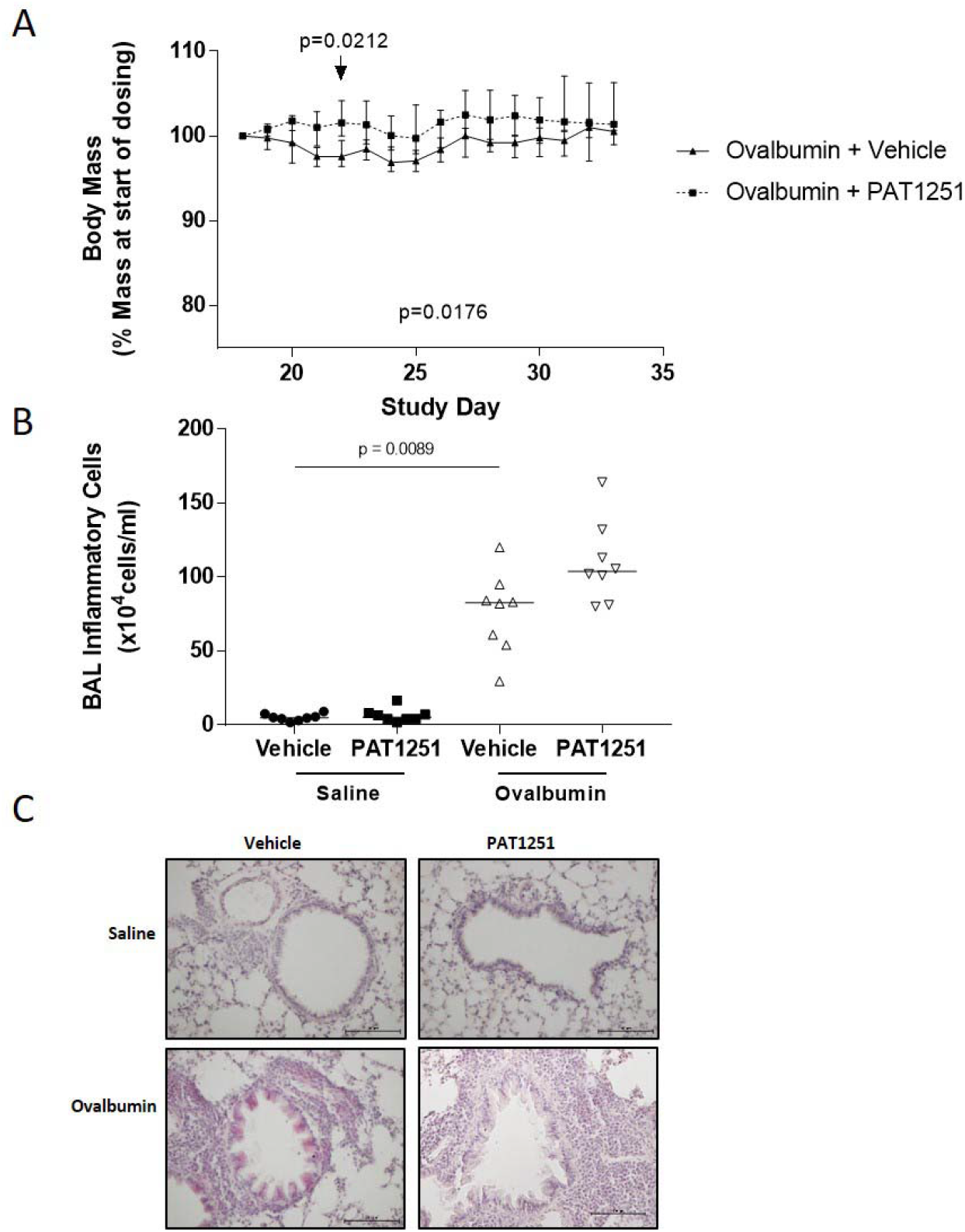
A Body mass data from mice repeatedly challenged with oropharyngeal ovalbumin that were treated daily with either vehicle control or LOXL2 Inhibitor, PAT1251 (30mg/kg). Data is shown from the day of the first drug dose i.e. 24 hours prior to the commencement of the ovalbumin challenges. The two groups were significantly different across time points (p=0.0176), and multiple comparison testing of individual time points gave p=0.0212 at day 22. Data are expressed as median ± interquartile range. Statistical analysis was performed using a 2-way ANOVA B Total number of inflammatory cells in BAL fluid was determined from animals subjected to ovalbumin model of asthma that were treated with either vehicle control or LOXL2 inhibitor (PAT1251; 30mg/kg). Data is expressed as median number of inflammatory cells. Statistical analysis was performed by non-parametric Kruskall Wallis test with Dunn’s multiple comparison test. C Representative images of haematoxylin and eosin stained lung tissue from animals subjected to ovalbumin model of asthma that were treated with either vehicle control or LOXL2 inhibitor (PAT1251; 30mg/kg). Images shown are at x20 magnification and are representative of n=8/group animals.

**Figure 7.**
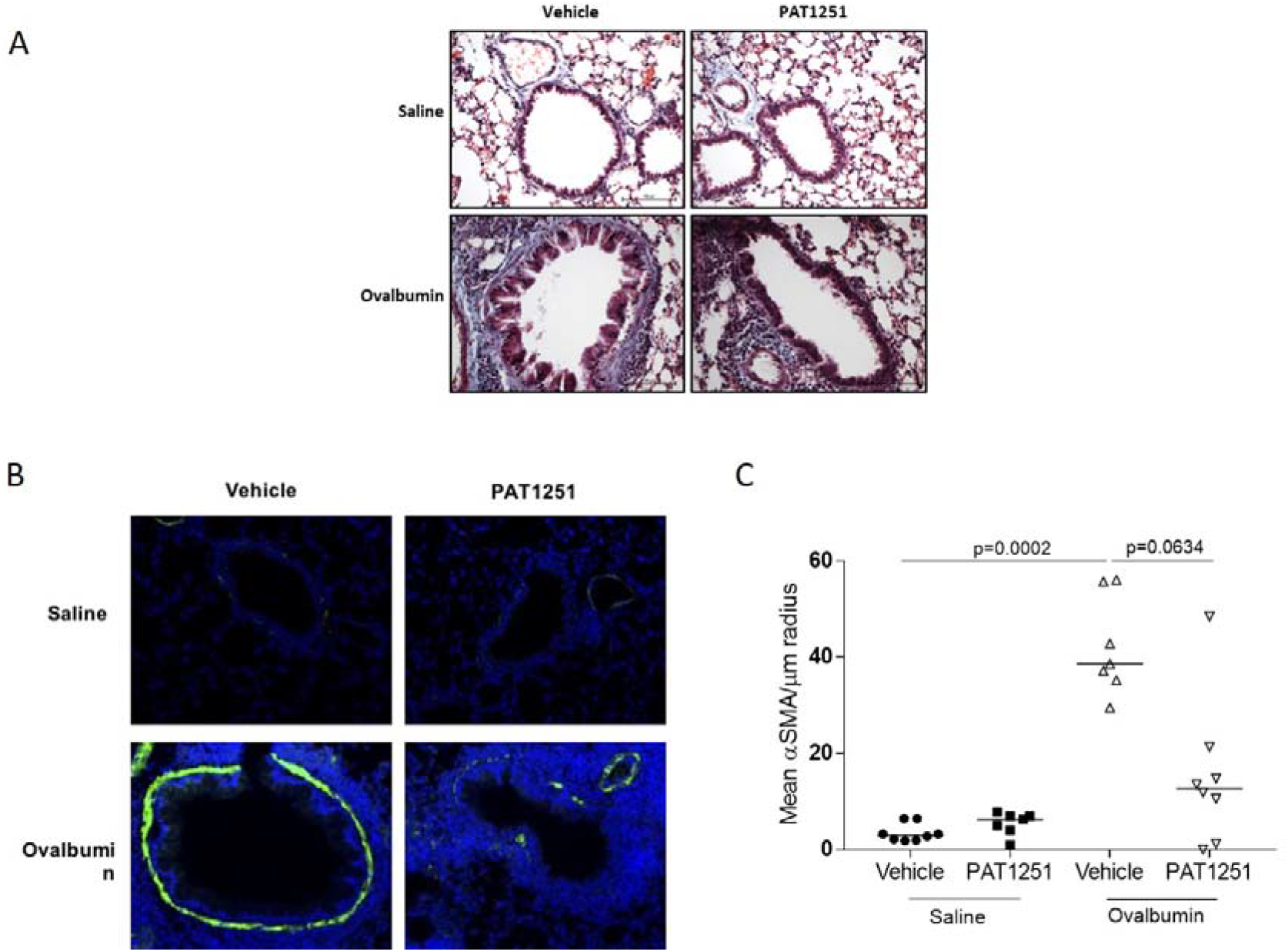
A Representative images of Masson’s trichrome stained lung tissue from animals subjected to ovalbumin model of asthma that were treated with either vehicle control or LOXL2 inhibitor (PAT1251; 30mg/kg). Images shown are at x20 magnification and are representative of n=8/group animals. B Representative immunofluorescent images of αSMA and Dapi stained lung tissue from animals subjected to ovalbumin model of asthma that were treated with either vehicle control or LOXL2 inhibitor (PAT1251; 30mg/kg). Images shown are at x20 magnification and are representative of the following group sizes: Saline + vehicle control n=8, saline + PAT1251 n=7, ovalbumin + vehicle n=7, ovalbumin + PAT1251 n=8. Representative negative control image is shown in Supplemental Figure S4F. C αSMA staining around airways less than 100μm in radius was quantified by measuring the area of the immunofluorescent staining. Data is expressed as median αSMA area per μm of airway radius. Statistical analysis was performed by non-parametric Kruskall Wallis test with Dunn’s multiple comparison test.

Ovalbumin challenge caused increased subepithelial collagen deposition compared with saline control challenged animals (Figure 6A). The amount of collagen surrounding airways in PAT1251 treated ovalbumin-challenged animals was reduced compared with ovalbumin-challenged vehicle controls (Figure 6A). Furthermore, ovalbumin challenge caused thickening of the ASM layer around small airways (<100μm radius) (Figure 6B). We demonstrated a greater than 2-fold decrease in the ASM thickness in PAT1251 treated animals compared with controls (Figure 6C).

## Discussion

Fibrogenesis and tissue remodelling is driven by ECM stiffness in multiple tissues and organs [7]. In severe asthma airway remodelling is associated with subepithelial fibrosis and thickening of the ASM layer, both of which may increase airway wall stiffness [12]. We have investigated how ECM influences TGFβ activation in ASM cells and identified a novel and important role for the matrix crosslinking enzyme LOXL2 in asthmatic airway remodelling.

TGFβ is a key driver of asthmatic airway remodelling and we have previously shown that ASM cells activate TGFβ in response to contractile agonists [21]. Here we demonstrate for the first time that basal TGFβ activation by ASM cells is affected by both ECM stiffness and by cyclical stretch confirming earlier studies in other cell types [30, 32]. Due to the confirmatory nature of these studies we used a smaller n than in subsequent experiments in order to preserve valuable primary human ASM cell cultures for hypothesis-driven exploratory experiments.

We found that asthmatic ASM cells are more contractile than non-asthmatic cells when measured using a collagen gel contraction assay and TGFβ activation correlates with contractility, confirming previously studies [33]. However, we found no difference in contractility between asthmatic and non-asthmatic cells when traction force microscopy (TFM) was utilised. This is potentially explained by the inherent differences the two assays. Collagen gels result in a 3D culture of ASM cells whereas TFM is performed on 2D monolayer. Crucially for our data interpretation, the weak correlation between TGFβ activation and collagen gel contractility combined with the TFM data, which more closely resembles the 2D nature of TMLC co-culture assays, led us to conclude that cell contractility differences are not responsible for increased TGFβ activation in asthmatic ASM cells.

It has long been postulated that the airway wall is stiffer in asthma, however, very few studies have directly investigated matrix stiffness in asthma. The passive stiffness of asthmatic smooth muscle explants is increased following length perturbations [34] and asthmatic fibroblasts have a higher Young’s elastic modulus than non-asthmatic controls [35]. This study is the first, to our knowledge, to directly measure the stiffness of ASM cell derived ECM and show that asthmatic ASM cells deposit stiffer ECM.

The ECM influences a variety of cellular pathways and can affect ASM cell biology [4, 5, 36–38] and impacts bronchoconstriction [39, 40]. The data presented here demonstrates a link between increased ASM cell ECM stiffness and asthmatic airway remodelling via TGFβ activation. While there is a paucity of data mechanistically linking increased ECM stiffness with airway remodelling, matrix stiffening activates pulmonary artery smooth muscle to trigger blood vessel remodelling [41, 42].

ECM stiffness is driven in part through crosslinking of matrix proteins. It has previously been reported that ASM cells can remodel extracellular collagen fibres [43]. LOXL2 is a crosslinking enzyme that has been investigated as a druggable target in fibrotic diseases [44]. Our data confirms an earlier study showing increased *LOXL2* gene expression in asthma [45] but also demonstrates that LOXL2 protein is increased in asthmatic ASM. LOXL2 is increased in several diseases associated with tissue remodelling [13–17, 44] but the functional consequences of increased LOXL2 in asthma have never been studied. We acknowledge that changes in LOXL2 expression do not necessarily lead to changes in enzymatic activity, however, there is currently no commercially available assay to measure LOXL2 activity. We therefore sought to investigate the functional effects of inhibiting LOXL2 activity in vitro and in vivo.

Our findings suggest that LOXL2 increases stiffness of ASM cell ECM, driving TGFβ activation and remodelling. Specifically, LOXL2 inhibition in vivo reduced allergen-induced airway remodelling. This is the first report linking LOXL2 with airway remodelling but previous studies link LOXL2 to vascular remodelling via effects on smooth muscle [46]. It is important to note that while animal models of fibrosis have shown promise for LOXL2 inhibition [17–20, 47] this has not yet translated to a clinical benefit in patients [48]. The reasons for this are unclear and there is merit in further consideration of LOXL2 inhibition in the treatment of asthmatic airway remodelling.

We have focused upon quantifying ASM layer thickness in smaller airways less than 100μm in radius in line with our previous work [21]. While airway remodelling changes are likely to affect airways of all sizes, we and others have found that the largest remodelling changes occur in small airways compared with large airways [21, 49, 50]. Furthermore, hyperpolarised gas magnetic resonance imaging of the lung has demonstrated significant regional heterogeneity in gas ventilation and distribution, suggestive of differential effects in smaller airways [51].

The data presented here highlight a potentially important role for LOXL2 in the development of asthmatic airway remodelling. We show that increased ECM can drive TGFβ activation in ASM cells and demonstrate that LOXL2 contributes to ECM stiffness. We demonstrate for the first time that LOXL2 expression is increased in asthma and we have shown that LOXL2 inhibition in vivo may reduce airway remodelling. Together our findings suggest that inhibition of LOXL2 may have merit as an approach to reduce airway remodelling in severe asthma.

## Acknowledgements

We would like to thank all patients and donors for consenting to provide samples for use in this research, and all support staff at the NIHR Biomedical Research Centres in Leicester and Nottingham for collecting and processing the samples.

This work was supported by a Medical Research Foundation/ Asthma UK personal fellowship and a NC3Rs David Sainsbury Fellowship, both held by A.L.T (MRFAUK-2015-312 and NC/K500501/1). J.R. received funding from the Newton Agham programme of the British Council (UK) and the Department of Science and Technology (Philippines). The human biopsy collection for use in LOXL2 immunohistochemistry was funded by a Medical Research Foundation grant (G1100163) held by S.R.J. Human ASM cells isolation was funded by the NIHR Biomedical Research Centres at Nottingham and Leicester.

## Supplemental Tables

**T1.**
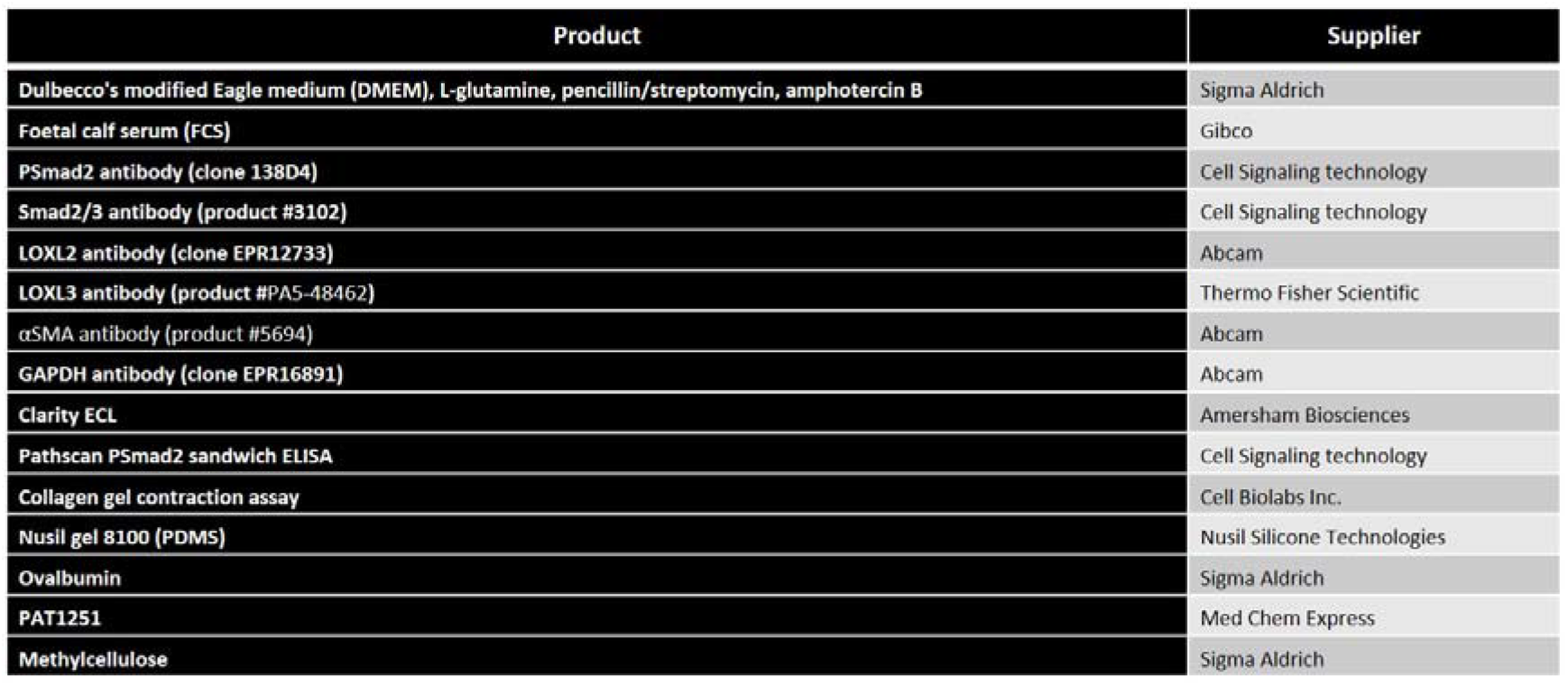
Table outlining further details of the materials used in the manuscript, including suppliers and clone details of all antibodies used.

**T2.**
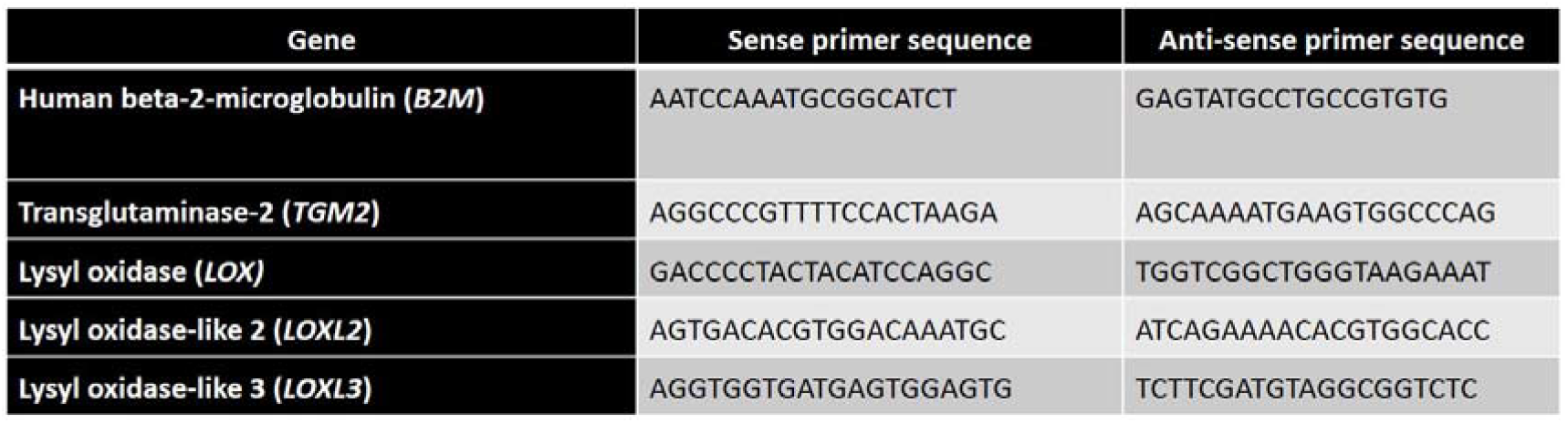
Table describing primer sequences for qRT-PCR. All primers were designed using Primer3 online software.

## Supplemental Figures

**S1.**
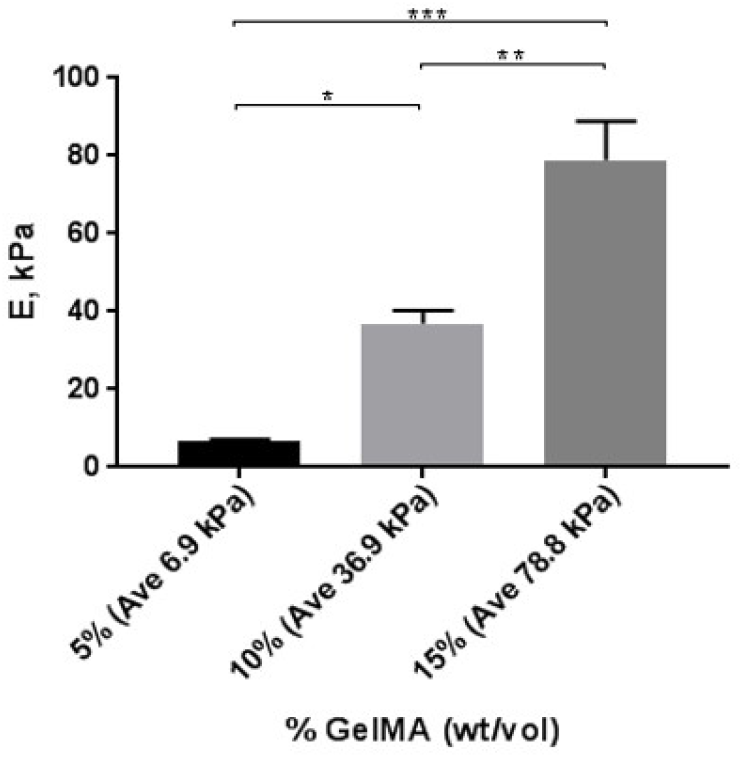
GelMa substrates (5, 10 and 15% GelMa wt/vol) were subjected to compressive modulus using texture analyser to determine the relative stiffness. 5% GelMa was approximately equivalent to the described stiffness of tracheal smooth muscle [22] and so was denoted as 1x stiffness in future experiments

**S2.**
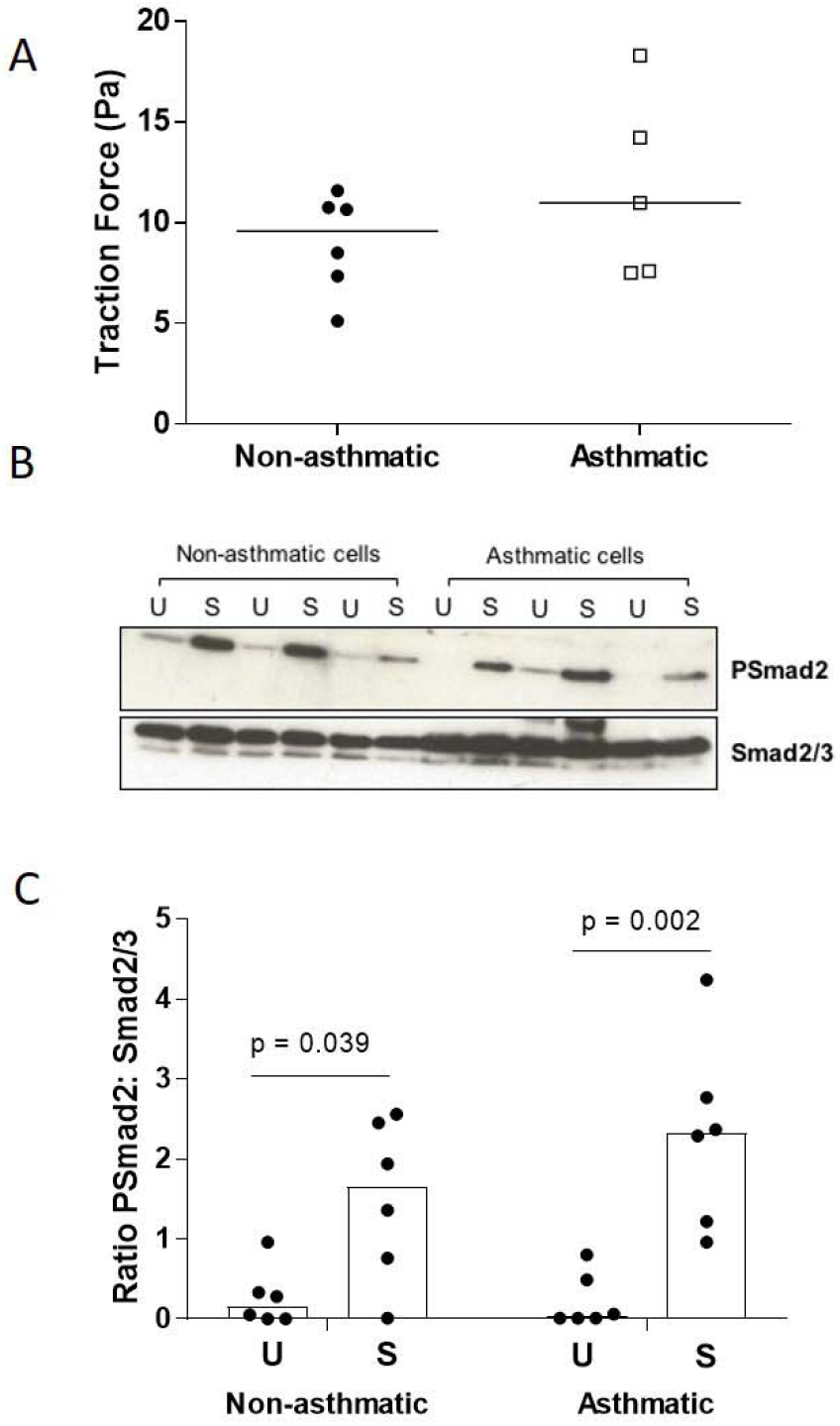
A Basal traction force imparted on a substrate by non-asthmatic and asthmatic ASM cells was determined using atomic force microscopy. Data is presented as median traction force (n) across all donor cell lines tested. Statistical analysis was performed using non-parametric Mann Whitney test. B Non-asthmatic and asthmatic ASM cells were cultured on Bioflex 6 well culture plates and either subjected to 15% stretch at 0.3Hz (S) or left unstretched (U) for four hours. PSmad2, total Smad2/3 and GAPDH were measured by western blotting. Figure shown is representative of n=6 non-asthmatic and n=6 asthmatic donor cell lines. C Densitometrical analysis of the western blots outlined in Figure S2B was performed using Image J and the data are shown here as median ratio of PSmad2: Smad2/3

**S3.**
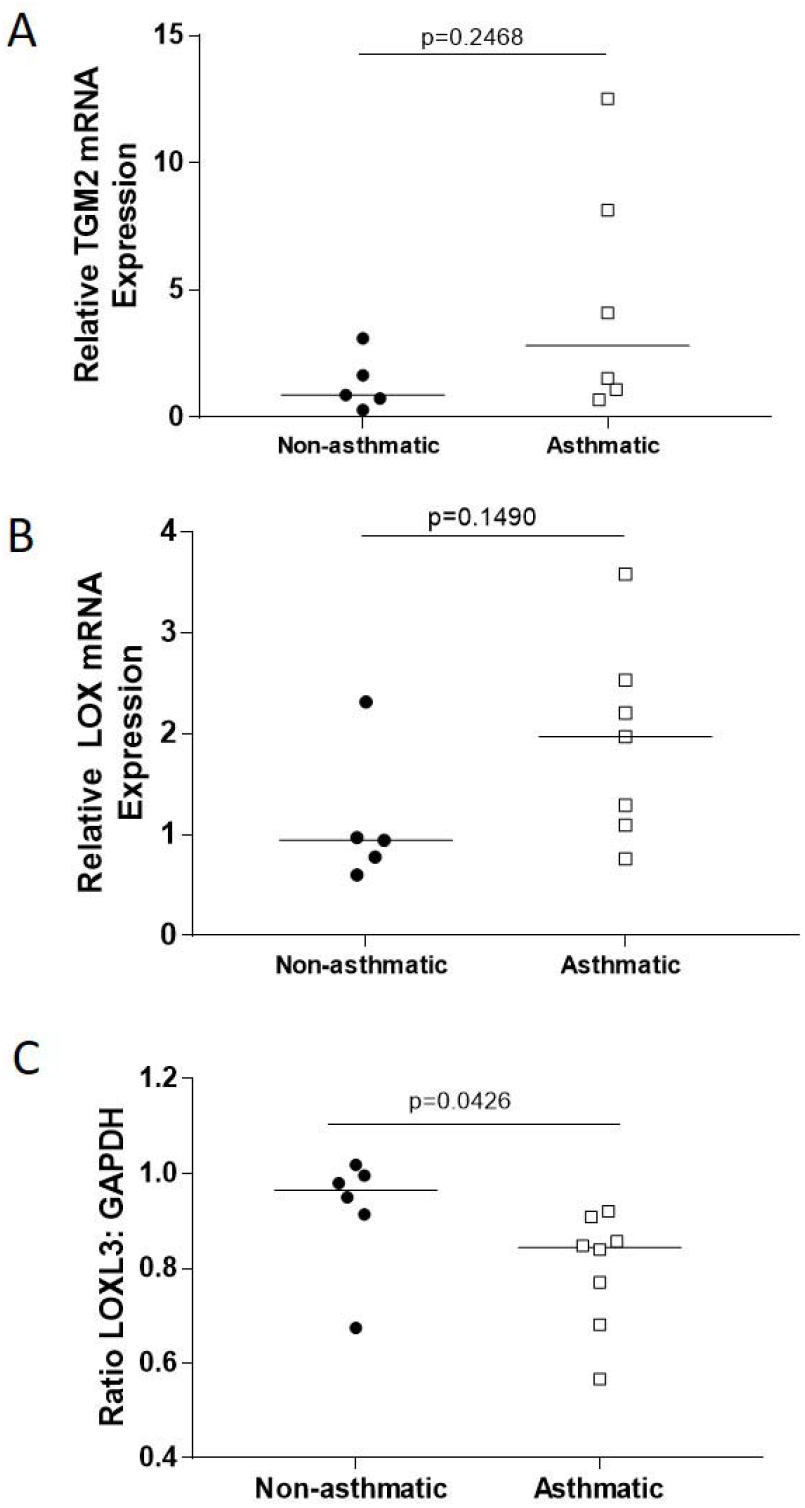
A Relative TGM2 mRNA in non-asthmatic and asthmatic ASM cells was determined by qRT-PCR. Data is expressed as median fold change in TGM2 mRNA versus the mean data from the non-asthmatic group. Statistical analysis performed by non-parametric Mann Whitney test. B Relative LOX mRNA in non-asthmatic and asthmatic ASM cells was determined by qRT-PCR. Data is expressed as median fold change in LOX mRNA versus the mean data from the non-asthmatic group. Statistical analysis performed by non-parametric Mann Whitney test. C Densitometrical analysis of western blots for LOXL3 (Figure 3C) in non-asthmatic and asthmatic ASM cells. Data is expressed as median ratio LOXL3 to GAPDH from two separate western blots (total n=6 non-asthmatic and n=8 asthmatic donor cell lines). Statistical analysis was performed by non-parametric Mann Whitney

**S4.**
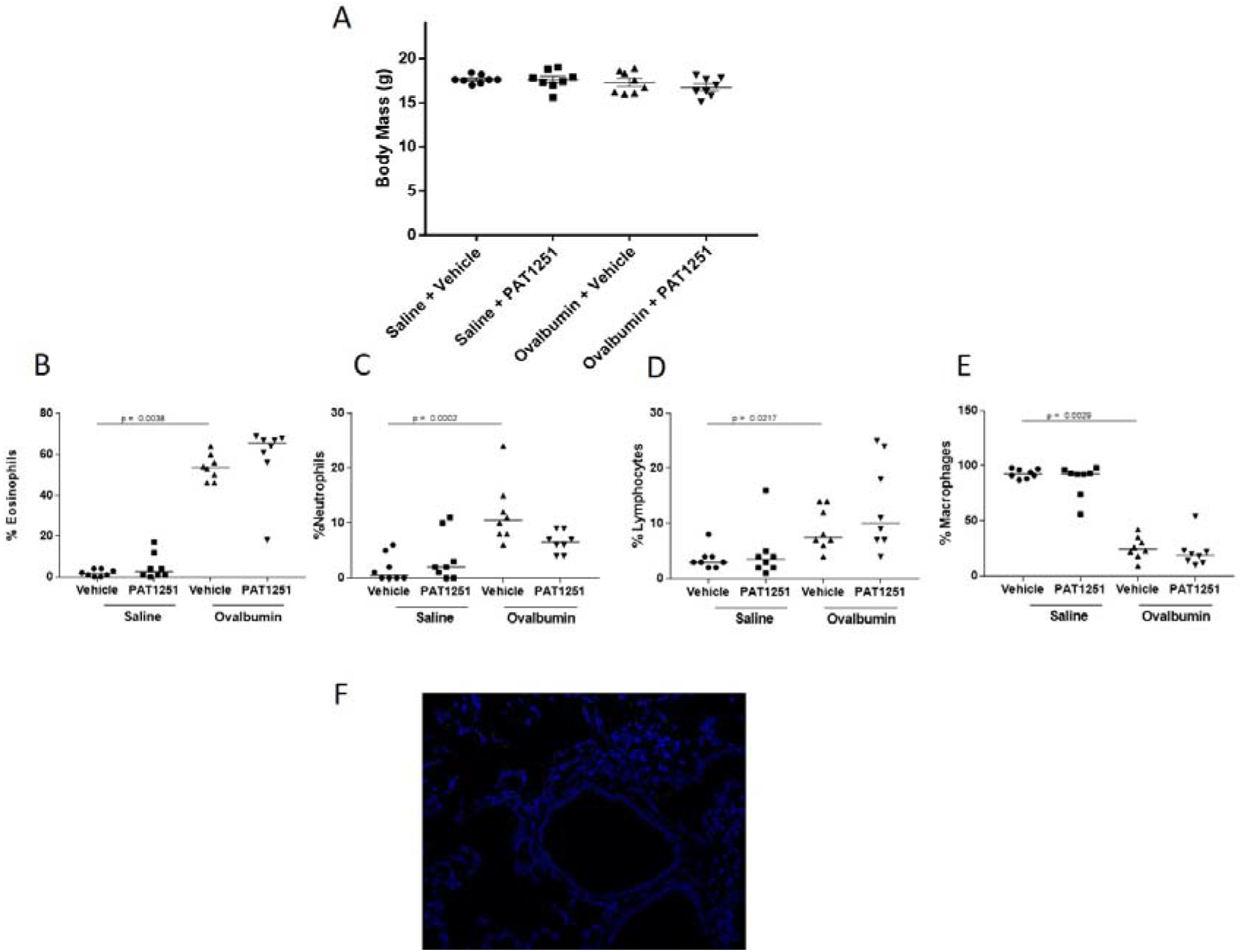
A Starting body mass data from mice allocated in to treatment groups in the in vivo model of asthma. Non-parametric Kruskall-Wallis with Dunn’s multiple comparison test showed there was no difference in starting mass between the four treatment groups. B Differential cell counts on BAL inflammatory cells were performed. The median percentage of eosinophils within the inflammatory cell population is expressed. Data were analysed by non-parametric Kruskall-Wallis with Dunn’s multiple comparison test C Differential cell counts on BAL inflammatory cells were performed. The median percentage of neutrophils within the inflammatory cell population is expressed. Data were analysed by non-parametric Kruskall-Wallis with Dunn’s multiple comparison test D Differential cell counts on BAL inflammatory cells were performed. The median percentage of lymphocytes within the inflammatory cell population is expressed. Data were analysed by non-parametric Kruskall-Wallis with Dunn’s multiple comparison test E Differential cell counts on BAL inflammatory cells were performed. The median percentage of macrophages within the inflammatory cell population is expressed. Data were analysed by non-parametric Kruskall-Wallis with Dunn’s multiple comparison test F Representative negative control image for αSMA immunofluorescence staining in Figure 7B.

## Supplemental Methods

### Cell culture

Human ASM cells were cultured from bronchial biopsies obtained at the Nottingham Respiratory Biomedical Research Unit, University of Nottingham (ethics number 08/H0407/1) and the Institute for Lung Health, University of Leicester (ethics number 08/H0406/189). Biopsies were obtained with written, informed consent from non-asthmatic and asthmatic donors. ASM cells were cultured from biopsy explants and used at passage 6. The cells were cultured in Dulbecco’s modified Eagles medium (DMEM) supplemented with 10% foetal bovine serum and 4mM L-glutamine at 37°C, 5% CO_2_ in a humidified incubator. Cells were growth arrested in serum-free DMEM for 24 hours prior to all experiments. A minimum of 3 donor cell lines were used in each experiment; in all experiments n is shown in the figures.

### Human Bronchial Biopsy Collection

Bronchial biopsies obtained from stable asthmatic and healthy control donors (n=6 per group) were used to determine the expression of LOXL2 protein within smooth muscle bundles. All tissue was collected at the Nottingham Biomedical Research Centre, University of Nottingham, under informed, written consent with ethical approval (12/EM/0199). Biopsies were obtained from 2^nd^ subdivision of right main bronchus. The asthmatic donors all had a formal diagnosis of asthma made by a medical practitioner, no history of exacerbation for 6 weeks prior to the bronchoscopy, and no recent use of oral corticosteroids, phosphodiesterase inhibitors or leukotriene inhibitors. Bronchial biopsies were fixed in formalin for 24 hours then paraffin wax embedded. The biopsies used have been previously published in [3]

### LOXL2 Inhibition in an In Vivo Ovalbumin Model

Studies were approved by the University of Nottingham Animal and Welfare Ethical Review Board (AWERB) and performed under Home Office personal, project and institutional license authority within the Animal (Scientific Procedures) Act 1986. Animals received free access to food (Tekland Global 18% protein rodent diet) and water, and were housed in specific pathogen free environment. 6-week-old female Balb/C mice (Charles River, UK) were sensitised to ovalbumin by intraperitoneal injection (i.p) of 10μg ovalbumin (Sigma Aldrich, UK) diluted 1:1 with Alum on days 0 and 12. Following sensitisation to ovalbumin animals were randomised to one of four treatment groups PBS + vehicle, PBS + PAT1251, OVA + vehicle, OVA + PAT1251. From day 18 animals received daily oral gavage of either 30mg/kg LOXL2 inhibitor (PAT1251; Med Chem Express, China) in 0.5% methylcellulose or 0.5% methylcellulose alone (100μl/dose). Animals were challenged with either 400 μg/ml ovalbumin in 50μl PBS or 50μl PBS alone via the oropharyngeal route under isoflurane anaesthesia on days 19, 20, 21, 22, 23, 24, 26, 28, 30 and 33. Animals were euthanised 24 hours after the final challenge. Bronchoalveolar lavage (BAL) wash performed using 1ml PBS. BAL inflammatory cells were cytospun (400rpm for 6 minutes) and stained with RapiDiff II kit (Atom Scientific, UK) according to the manufacturer’s instructions. A differential cell count of eosinophils, monocytes, neutrophils and leukocytes was performed using a Nikon Eclipse 90i microscope. Total BAL inflammatory cell count was performed using a haemocytometer.

### Statistical Analysis

All statistical tests were discussed with and approved by a statistician (I.D.S). Data are reported as a median of n observations and non-parametric tests were chosen due to the relatively small sample sizes restricting assumptions of a normal distribution and so that tests were not affected by outliers. Mann-Whitney or Kruskall-Wallis tests were used to compare two, or more than two groups, respectively. All ECM crossover experiments were analysed using a one-sample t-test versus the effect on the cells’ own ECM. Details of the specific statistical test used for each figure is included in the figure legend. In all in vivo studies the experimental unit (n) denotes an individual animal. All analysis was performed using Graphpad Prism (v7.04, La Jolla, USA)

